# Single cell transcriptional landscape of long non-coding RNAs orchestrating mouse heart development

**DOI:** 10.1101/2022.04.29.490042

**Authors:** Thaís A. R. Ramos, Soo Young Kim, Thomas G. Gillette, Joseph A. Hill, Sergio Lavandero, Thaís G. do Rêgo, Vinicius Maracaja-Coutinho

## Abstract

Long non-coding RNAs (lncRNAs) comprise the most representative transcriptional units of the mammalian genome, and they’re associated with organ development that can be associated with the emergence of diseases, such as cardiovascular diseases. Thus, we used bioinformatic approaches, machine learning algorithms and statistical techniques to define lncRNAs involved in mammalian cardiac development. We used a single-cell transcriptome dataset generated from 4 embryonic and 4 postnatal stages. Our study identified 8 distinct cell types, novel marker transcripts (coding/lncRNAs) and also, differential expression and functional enrichment analysis reveal cardiomyocyte subpopulations associated with cardiac function; meanwhile modular co-expression analysis reveals cell-specific functional insights for lncRNAs during myocardial development, including a potential association with key genes related to disease and the “fetal gene program”. Our results evidence the role of particular lncRNAs in heart development, and highlights the usage of co-expression modular approaches in the cell-type functional definition.

## Introduction

Approximately 90% of the mammalian genome is transcribed in a cell-specific manner into RNA [1], but only 2-3% is transcribed to protein-coding messenger RNA (mRNA) [2,3]. The vast majority of RNA transcripts are grouped into non-coding RNAs (ncRNAs), which are now routinely investigated in almost every transcriptome or genome project. Non-coding RNAs can be classified according to their functional role (*i*.*e*. regulation, translation, splicing) and can be divided into two main classes according to their length: small non-coding RNAs (<200 nucleotides (nt)) and long non-coding RNAs (lncRNAs) (≥200□nt) [2–4].

Long non-coding RNAs (lncRNAs) comprise the most representative transcriptional units of the mammalian genome and are associated with organ development. A large number of lncRNAs are generated by RNA polymerase II as transcripts “similar” to mRNAs. They can also be subject to splicing and polyadenylation, but they do not have the potential to encode proteins [5]. LncRNAs can also control the flux of genetic information through various gene regulation mechanisms, including transcriptional interference, chromatin modifications, mRNA stability, modulation of alternative splicing and chromosome structure, and post-translational modifications [5,6]. In summary, lncRNAs, in interaction with other molecules, coordinate multiple physiological processes.

LncRNAs are widely recognized to have a profound impact on fine-tuning the regulation of multiple critical biological processes, exhibiting cell type- and tissue-specific expression. Recently emerging evidence has demonstrated that lncRNAs can determine the reprogramming of lineage fates and cell subtypes [7,8]. However, little is known regarding the expression and function of lncRNAs in heart development [9]. In addition, it is unclear whether lncRNAs are highly expressed in subsets of cells within the tissues. The single-cell expression patterns of lncRNAs may help uncover novel biological functions for different cell populations. Due to its role in the fine-tuning regulation of crucial cellular biological processes, measuring the abundances of ncRNAs in individual cells is critical to defining cell-type particularities and single-cell functions [10].

Recently, a new type of lncRNA has been identified based on the evolutionary conservation of their promoters and their syntenic positions between humans and mice, the so-called positionally conserved lncRNAs (pcRNAs) [6]. These lncRNAs are located in regions enriched with neighboring genes that could play essential roles in developing cardiovascular tissue, arterial morphogenesis, and smooth muscle tissue cell differentiation. Furthermore, tissue-specific expression patterns shared between humans and mice revealed a group of pcRNAs with exclusive expression in the heart. Furthermore, a subset of these transcripts manifested potential importance in chromatin organization based on its localization that overlaps the binding sites for the CTCF chromatin organizer, the chromatin loop anchor points, and borders of topologically associating domains [6].

Mutations and misregulation of ncRNAs have been shown to contribute to multiple disease processes, such as cardiovascular disease, cancer, diabetes, and cerebral ischemic stroke [3,6]. The involvement of ncRNAs in cardiovascular diseases has been well described [11,12]. LncRNAs have been associated with many disorders and some of them have been identified as key regulators in the development and progression of cardiovascular diseases [3]. Strikingly, the failing adult heart resembles the fetal heart in many ways. Several gene expression changes that have been reported in failing hearts are consistent with the ‘re-expression of the fetal gene expression program’ [13,14]. In an effort to develop novel diagnostic and therapeutic strategies for heart disease, it is essential to define the molecular elements active in developmental heart stages (from embryo to adulthood). Dissecting patterns and cell-specific distributions of coding and non-coding RNAs during development will enable us to unveil novel regulatory processes occurring during cardiac development and potentially lead to improved therapeutic targeting [2–4].

Our work reported here employed a single-cell RNA sequencing (scRNA-seq) dataset to identify and quantify lncRNAs in multiple cardiac cell types during the course of murine heart development. To this end, we used a combination of machine learning and statistical techniques to identify the markers and assign the cell types. Systems biology analyses identified co-expression modules linked to heart development and cardiovascular diseases. We also uncovered differentially expressed transcripts in subpopulations of cardiomyocytes. Finally, from this work we were able to identify eight cardiac cell types; several new coding, lncRNA, and pcRNA markers; 2 cardiomyocyte subpopulations at 4 different time points (ventricle E9.5, left ventricle E11.5, right ventricle E14.5 and left atrium P0) that harbored co-expressed gene modules enriched in mitochondrial, heart development and cardiovascular diseases.

## Results

### Expression quantification and cell-type classification according to expression patterns

The single-cell transcriptomic dataset used in the study included data from four embryonic (E9.5, E11.5, E14.5, E18.5) and four postnatal (P0, P3, P7, P21) stages from five distinct myocardial anatomical areas: ventricle, atrium, left ventricle, right ventricle and left atrium (Fig. 1). The left ventricle was the only compartment with data collected across all time points. It is worth mentioning that at embryonic stage 9.5 (E9.5) the atrium and ventricle are not yet separated as left and right compartments. We were able to estimate the expression of dozens of coding and long non-coding transcripts identified in specific compartments and time points (Table 1). For lncRNAs, we found from 738 (left ventricle P21) to 2,636 (left ventricle P7) transcripts, with only P21 with fewer than 1,000 lncRNAs. These observations are in accordance with the number of lncRNAs reported in the literature dissecting the non-coding transcriptome with a single-cell focus [15,16].

**Fig. 1.**
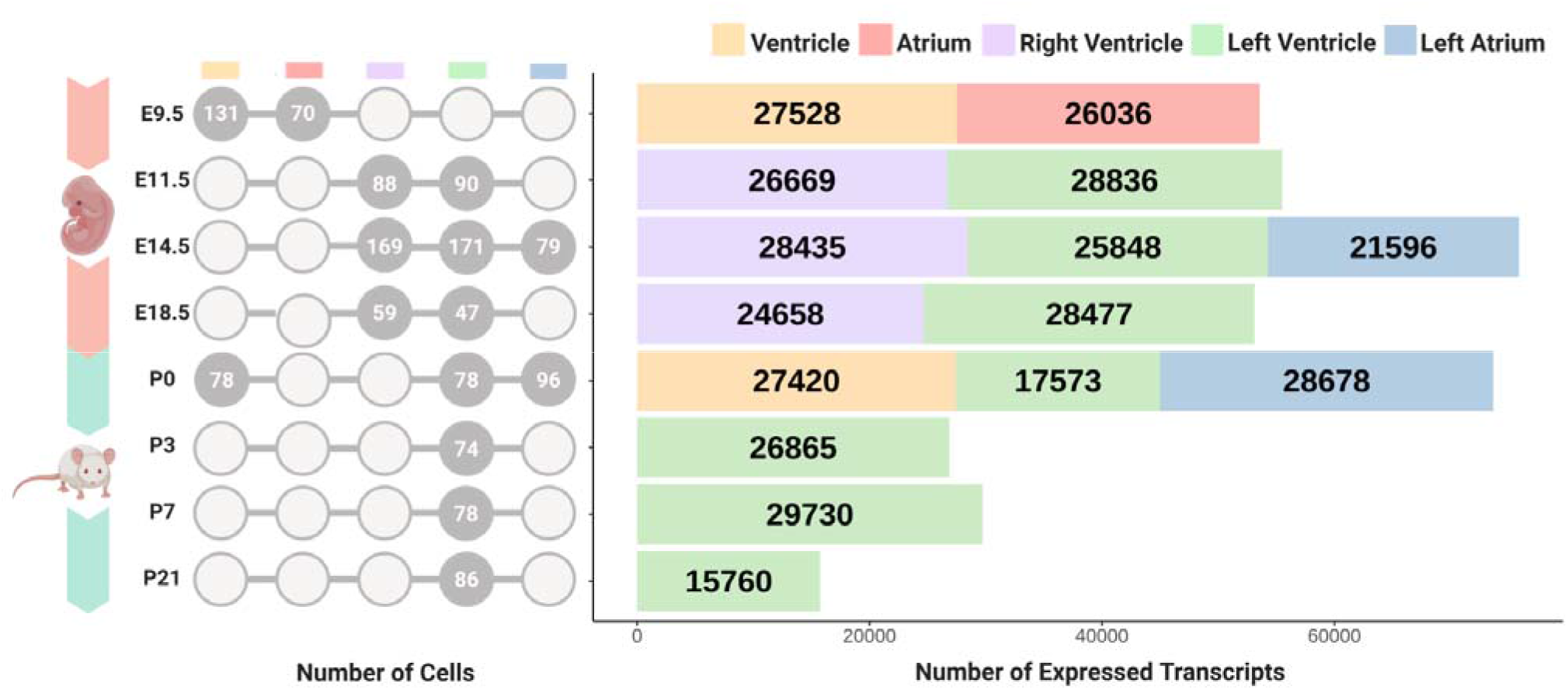
Single-cell transcriptomics dataset based on the number of cells and transcripts in each myocardial compartment across developmental stages. Circular diagram reports the number of single cells for each data point. The bar graph shows the number of expressed transcripts color-coded by anatomical area (yellow: ventricle, red: atrium, purple: right ventricle, green: left ventricle, blue: left atrium).

**Table 1.** Number of transcripts expressed in Fig. 1, marker transcripts (coding genes, lncRNAs and pcRNAs) and distribution in each compartment according to development stage. Expressed transcripts and marker transcripts are reported according to each of the three classes (coding, lncRNAs, and pcRNAs) in each compartment across developmental stages. The first column is the development stage; the second column represents the compartment; 3rd and 4th columns are related to the number of expressed coding genes and markers coding genes, respectively; 5th and 6th columns report expressed lncRNAs and markers lncRNAs, respectively; finally, 7th and 8th columns reported expressed pcRNAs and markers pcRNAs, respectively.

| Stage | Compartment | Coding genes |  | lncRNAs |  | pcRNAs |  |
| --- | --- | --- | --- | --- | --- | --- | --- |
|  |  | Total | Markers | Total | Markers | Total | Markers |
| **E9.5** | Ventricle | 25,217 | 1,770 | 2,190 | 186 | 121 | 13 |
| **E9.5** | Atrium | 24,181 | 453 | 1,751 | 27 | 104 | 4 |
| **E11.5** | Right ventricle | 25,056 | 1,432 | 1,514 | 96 | 99 | 11 |
| **E11.5** | Left ventricle | 26,468 | 1,767 | 2,246 | 241 | 122 | 11 |
| **E14.5** | Right ventricle | 25,826 | 2,939 | 2,467 | 448 | 142 | 24 |
| **E14.5** | Left ventricle | 23,929 | 1,446 | 1,814 | 154 | 105 | 14 |
| **E14.5** | Left atrium | 20,077 | 329 | 1,438 | 17 | 81 | 3 |
| **E18.5** | Right ventricle | 22,771 | 221 | 1,786 | 24 | 101 | 0 |
| **E18.5** | Left ventricle | 26,150 | 387 | 2,194 | 38 | 133 | 5 |
| **P0** | Ventricle | 25,428 | 673 | 1,880 | 70 | 112 | 6 |
| **P0** | Left ventricle | 16,453 | 63 | 1,064 | 7 | 56 | 2 |
| **P0** | Left atrium | 26,350 | 757 | 2,194 | 122 | 134 | 10 |
| **P3** | Left ventricle | 24,582 | 632 | 2,174 | 68 | 109 | 2 |
| **P7** | Left ventricle | 26,957 | 574 | 2,636 | 60 | 137 | 2 |
| **P21** | Left ventricle | 14,984 | 67 | 738 | 6 | 38 | 0 |

Next, we used the Silhouette method with hierarchical clustering to identify the optimal number of clusters in each sample, combined with the chi-squared and adherence tests to determine statistical significance (*i*.*e*. p-value<0.05) of each cluster and make cell-type assignments. t-SNE plots were used for cluster visualization, manifesting how cell types are distributed in space according to transcript expression patterns and cell type (Fig. 2). Heatmaps (Supplementary Fig. 1) demonstrate transcript expression patterns according to cell type and corroborate the t-SNE cell subtype assignments (Fig. 2).

**Fig. 2.**
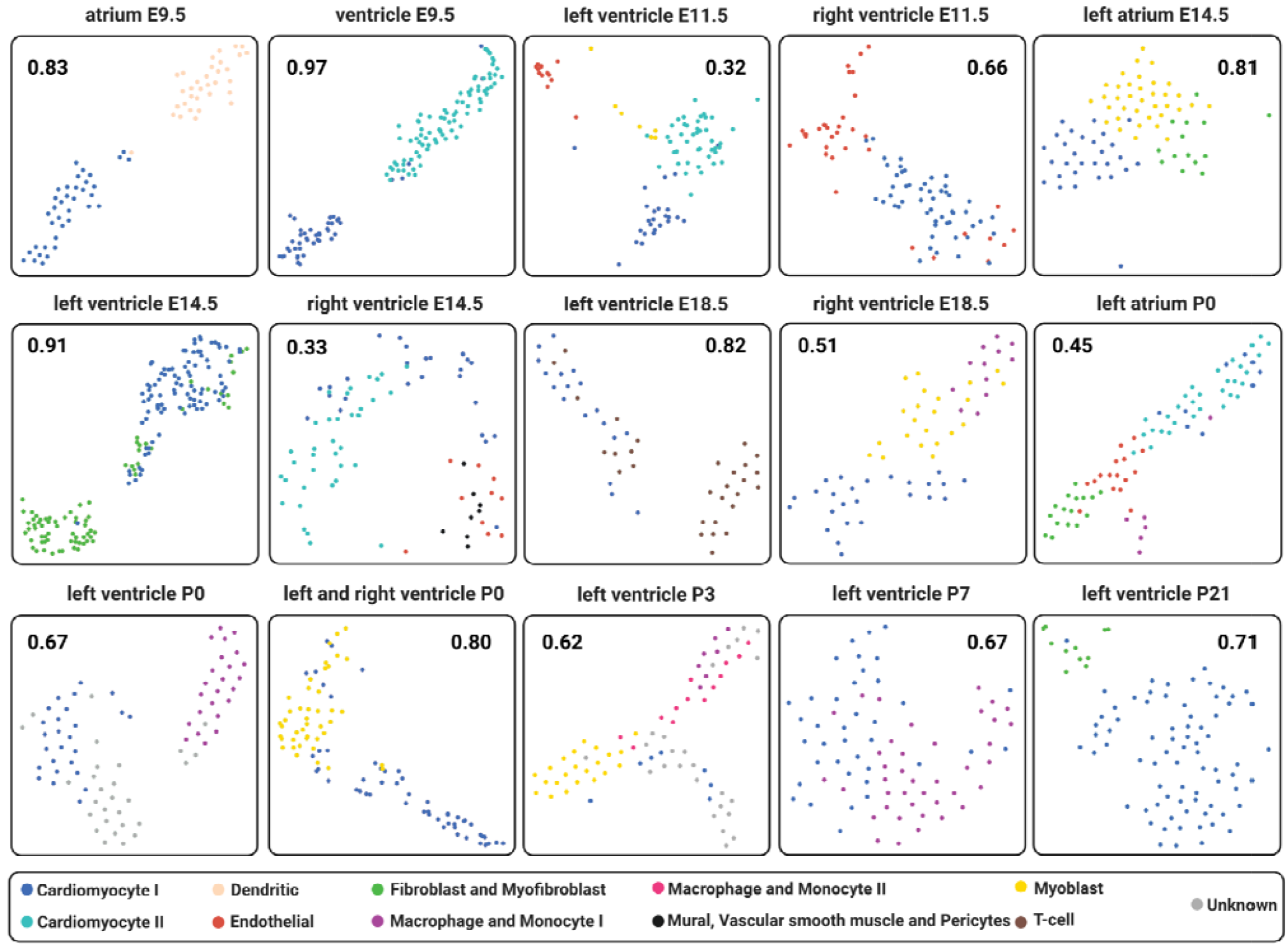
Representative cell-type assignment in each anatomical area per time point. t-SNE plots in which each square represents a compartment at a specific development stage. Each dot represents a cell, and each color describes a particular cell type: Blue - Cardiomyocyte I; Light blue - Cardiomyocyte II; Salmon - Dendritic; Red - Endothelial; Green - Fibroblast and Myofibroblast; Purple - Macrophage and Monocyte II; Black - Mural, Vascular smooth muscle and Pericytes; Myoblast - Yellow; Brown - T-cell; and Gray - Unknown cell types (*i*.*e*. can be a different cell type in which we don’t have markers). In addition, each number inside the squares is related to the silhouette coefficient score for each compartment and time point.

We obtained a maximum of 5 potential cell clusters in the postnatal development stages (*e*.*g*. left atrium P0 and left ventricle P3) with 4 clusters in the ventricle compartment in embryonic stages (*e*.*g*. left ventricle E11.5 and right ventricle E14.5). We observed unique compartments and developmental stages with 3 (*e*.*g*. left atrium E14.5, right ventricle E18.5 and left ventricle P0) and 2 clusters (the other 8 compartments and stages). The list of potential gene markers identified based on their expression patterns is listed in Supplementary File 1.

To classify clusters by cell type, we generated a comprehensive cell marker database by combining information retrieved from three datasets available in the literature [17–19] and the PanglaoDB database [20]. We were able to assign 8 distinct cell types in all compartments and time points: cardiomyocytes, myoblasts, endothelial cells, vascular smooth muscle cells and pericytes, fibroblasts and myofibroblasts, T-cells, macrophages and monocytes, and dendritic cells (t-SNE plots in Fig. 2). Moreover, we identified two subpopulations of cardiomyocytes in four compartments and time-points (ventricle E9.5, left ventricle E11.5, right ventricle E14.5, and left atrium P0), as well as two macrophage and monocyte subpopulations in left ventricle P3. A list with all markers identified for each cell type through each stage of heart development is presented (Supplementary File 1). The heatmaps and dendrograms of the hierarchical clustering for each heart chamber and time-point, corroborating the subtype assignment, can be accessed in Supplementary Fig. 1.

### Single-cell transcriptomics reveal lncRNAs as potential cell markers for eight cell types during heart development

After identifying and assigning cell types, we analyzed changes in transcripts in each cell type per developmental stage. Differentially expressed transcripts were divided into 3 classes: coding transcripts, lncRNAs, and pcRNAs (Fig. 3, Supplementary File 1). We identified at least one lncRNA (cardiomyocytes, left atrium E14.5) as a gene marker in all time points and chambers evaluated, reaching up to 197 in cardiomyocytes in the right ventricle E14.5. Positionally conserved non-coding RNAs were classified as markers for several cell types. As expected, our data show that cardiomyocyte populations are present in all compartments in all development stages [17,21]. They also reveal specific cell populations per region and developmental stage. For instance, a cell population with a dendritic cell-like transcriptional signature was observed exclusively in E9.5 atrium; meanwhile, we identified a vascular smooth muscle-like population in E14.5 right ventricle and a T cell-like population in E18.5 left ventricle. Interestingly, fibroblasts and myofibroblast transcriptional signatures were detected in E14.5, a stage at which fibroblasts from the epicardium are known to undergo epithelial-to-mesenchymal transition (EMT) [22]. Cell populations manifesting monocyte and macrophage profiles appeared only at late embryonic and postnatal stages, matching previously reported evidence of monocyte-derived macrophages increasing in the myocardium post-birth [23].

**Fig 3.**
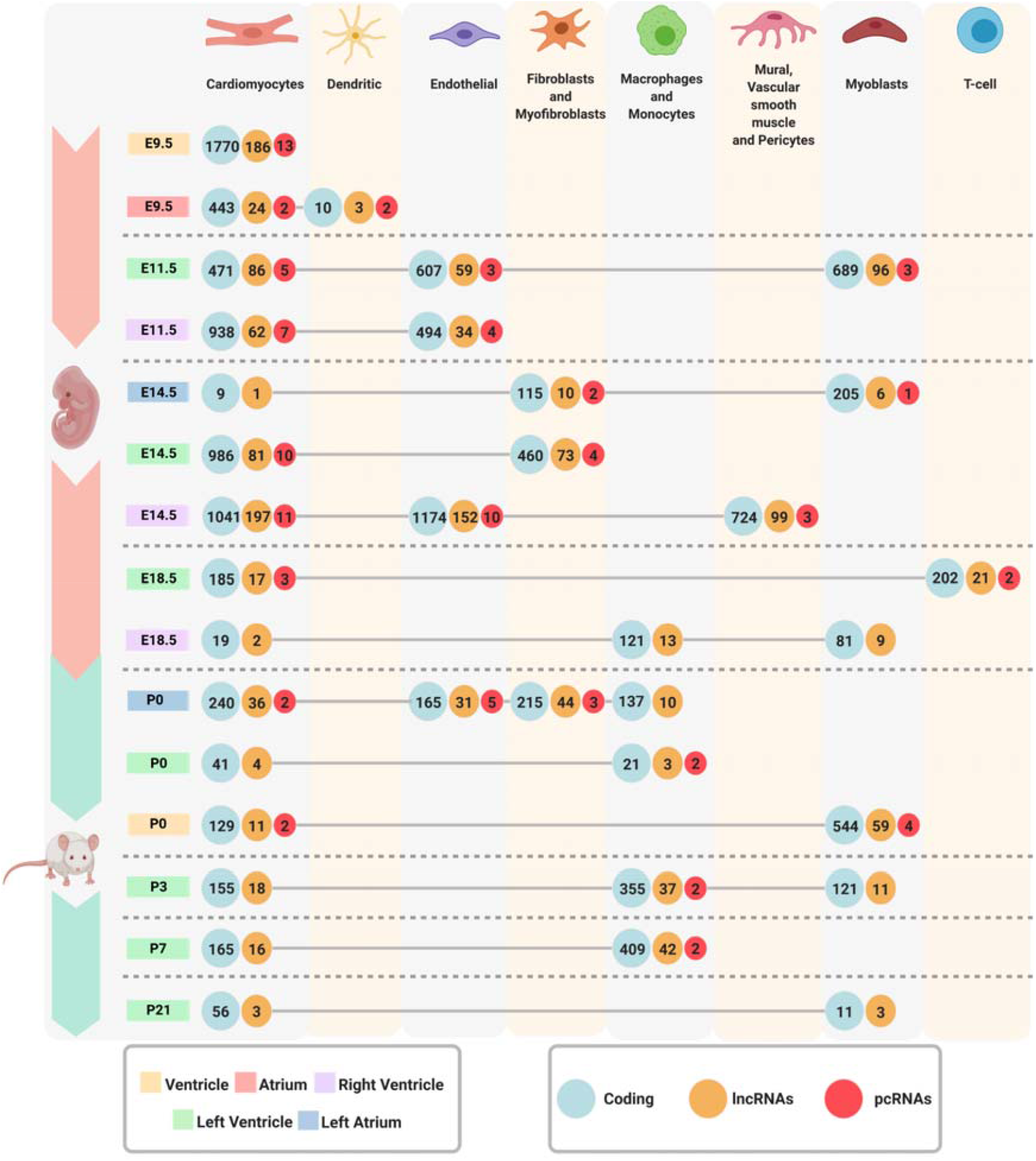
Cell marker transcripts by cell type and chamber during heart development. The numbers inside each color-coded circle (green: coding genes, orange: lncRNAs, red: pcRNAs) represent the number of marker’s transcripts. Each compartment is represented by a color: Yellow - Ventricle; Red - Atrium; Purple - Right Ventricle; Green - Left Ventricle; and Blue - Left Atrium. In addition, we found marker’s through 8 different cell types represented by each column.

In addition, we tested the cell types that can be identified in the same anatomical region (*e*.*g*. fibroblasts and myofibroblasts in the left atrium [24]). At the E9.5, non-fibroblast-enriched cells were identified in atrium or ventricle compartments, consistent with a prior report [17]. We argue that our cell type analysis based on coding and lncRNA profiles may reflect unique aspects of myocardial developmental events. In addition, it not only matches analysis based on the coding transcriptome alone [17] but also reveals new cell populations not uncovered in that analysis (Fig. 3).

### scRNA-seq identifies specific signatures for coding genes and lncRNAs for the same cell type at different heart compartments

Next, we compared the expression profile of cell-type specific transcript markers between the different cardiac chambers. We evaluated the presence/absence of each marker of the 4 most abundant cell populations (cardiomyocytes, endothelial cells, fibroblasts, macrophages and monocytes) per time during the course of myocardial development (Fig. 4). Interestingly, our analysis revealed multiple unique coding and non-coding RNA (lncRNAs, pcRNAs) signatures for the same cell type in different chambers, suggesting the same cell type adopts novel compartment-specific gene expression patterns per anatomical milieu. Overall, coding and non-coding profiles were similar in terms of proportions of shared and unique transcripts across anatomical areas at all time points, suggesting that one type of transcript is neither more nor less specific for the anatomical origin.

**Fig. 4.**
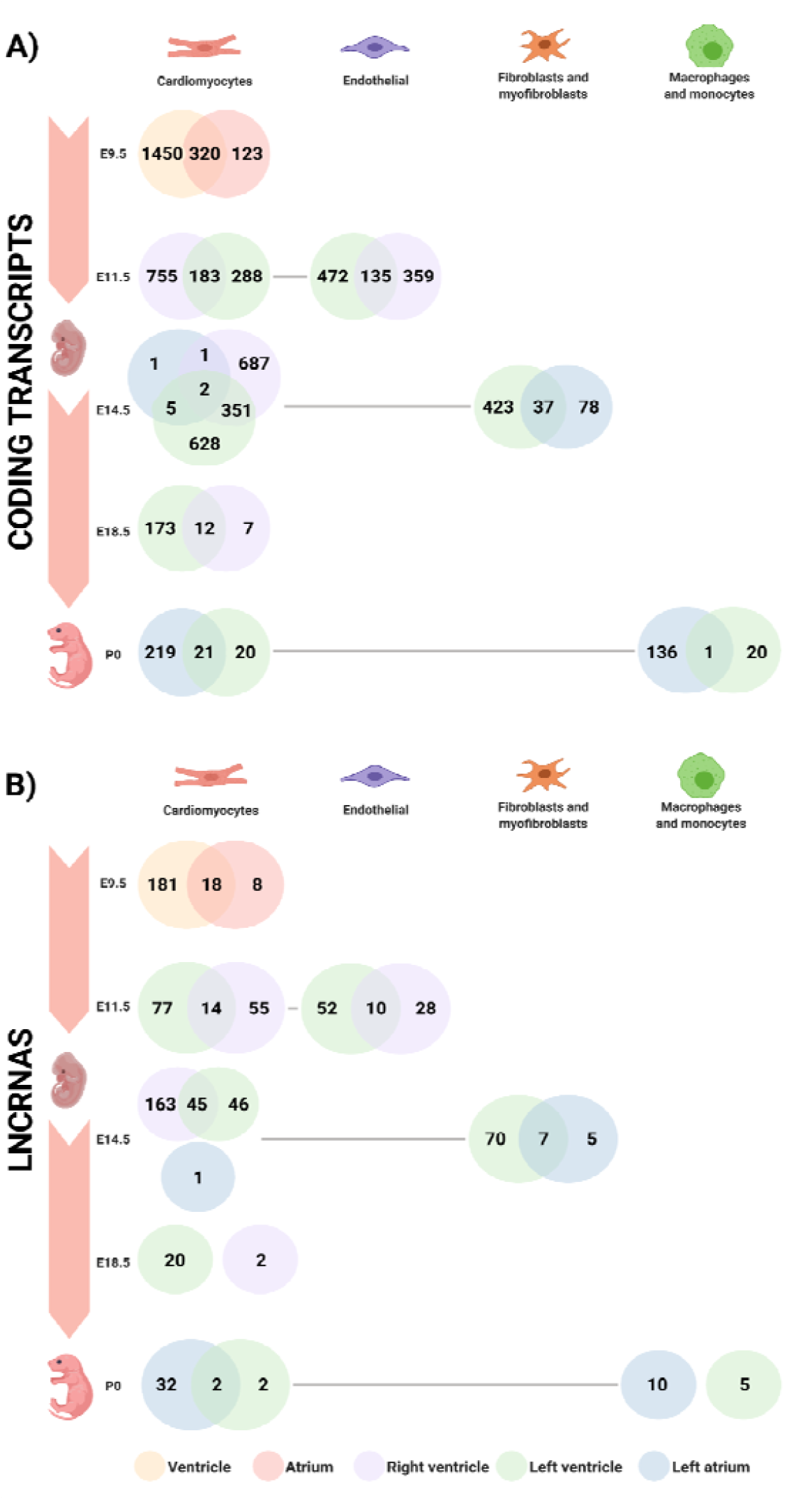
Venn diagram depicting differentially expressed transcripts across cardiac chambers during development. Venn diagrams are color coded to represent a compartment of the heart: Yellow - Ventricle; Red - Atrium; Purple - Right Ventricle; Green - Left Ventricle; and Blue - Left Atrium. The diagram was created from E9.5 to P0, since P3, P7 and P21 only had left ventricle cells; thus, the compartment’s comparisons were not possible in these 3-time points. The number of transcripts is noted in the diagram through 4 different cell types (Cardiomyocytes; Endothelial; Fibroblasts and Myofibroblasts; Macrophages and Monocytes) represented by each column. **A)** This analysis shows the number of shared and unique coding transcripts of each compartment. **B)** This analysis shows the number of shared and unique non-coding transcripts (lncRNA and pcRNA) of each compartment.

### Differential expression and functional enrichment analysis reveal cardiomyocyte subpopulations associated with metabolic diseases, mitochondria, and cardiac function

To pursue differences within the identified cardiomyocyte subpopulations, we analyzed differential expression to characterize biological variation between the 2 subpopulations (Cardiomyocyte 1 and 2) identified in each chamber and developmental stage (ventricle E9.5, left ventricle E11.5, right ventricle E14.5, and left atrium P0). Different cardiomyocyte cell types were identified in distinct chambers and time points (Fig. 5), with the ventricle population of cells presenting greater number of differentially expressed genes across both cardiomyocyte types, with 3,568 coding and 244 non-coding transcripts detected as differentially expressed at embryonic stage E9.5 (ventricle); 806 coding and 53 non-coding transcripts in the embryonic stage E11.5 (left ventricle); and other 3,173 coding and 222 non-coding transcripts in the embryonic stage E14.5 (right ventricle). On the other hand, we identified distinct cardiomyocyte sub-populations in the left atrium at the post-natal time point P0, with 84 coding and 7 non-coding transcripts identified as differentially expressed (Supplementary File 2).

**Fig. 5.**
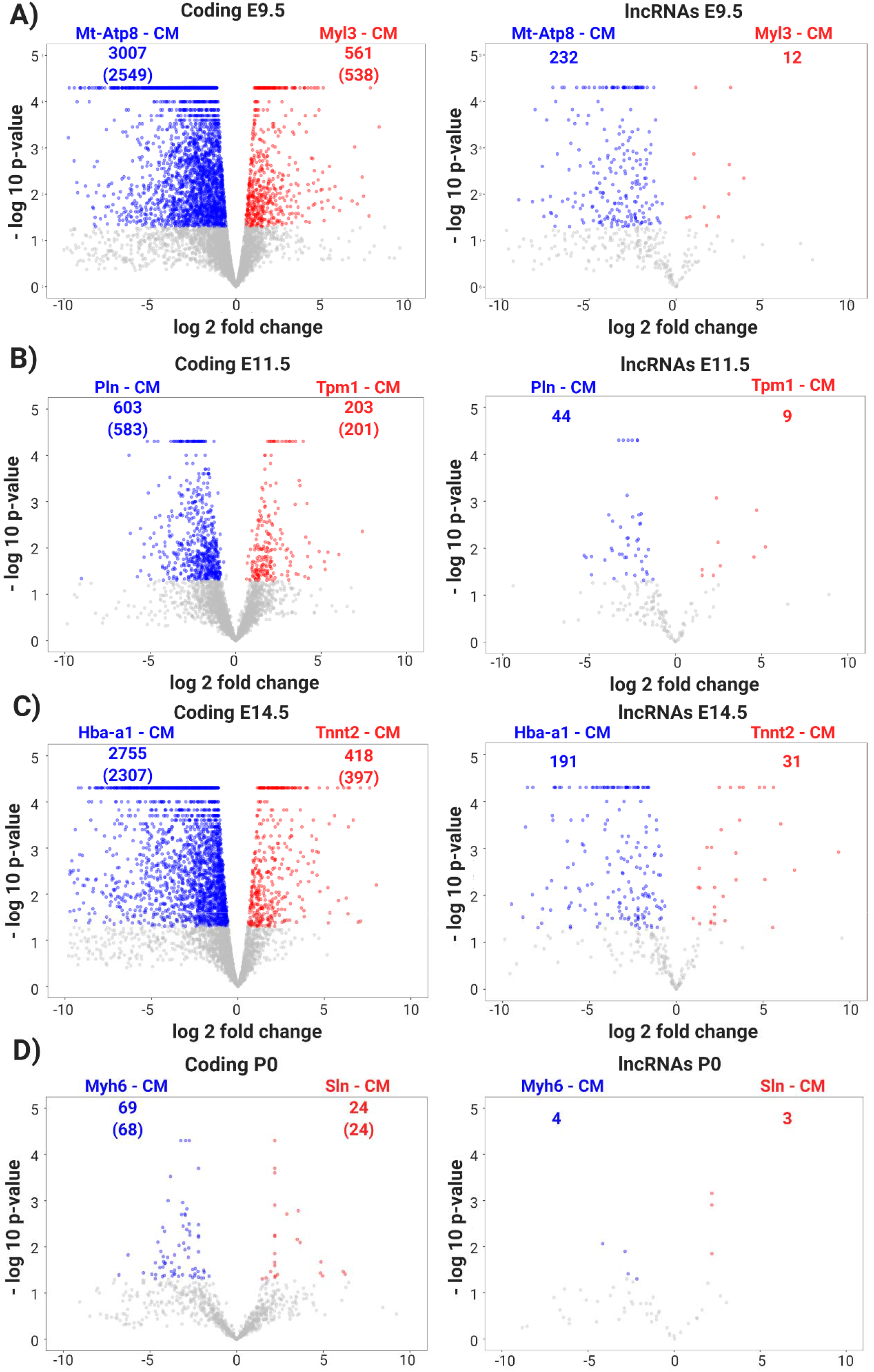
Volcano plots depicting cardiomyocyte subpopulations during the course of heart development. The left side (**A, C, E, G**) are related to coding transcripts in ventricle E9.5, left ventricle E11.5, right ventricle E14.5 and left atrium P0, respectively. The right side (**B, D, F, H**) are related to lncRNAs transcripts in ventricle E9.5, left ventricle E11.5, right ventricle E14.5 and left atrium P0, respectively. The blue and red colors are related to down- and up-regulated transcripts, respectively; the numbers are related to differentially expressed transcripts in each condition (down- and up-regulated); and the numbers in parenthesis are the number of differentially expressed genes.

We named the cardiomyocyte sub-populations according to the coding gene with the highest expression for each chamber and time point. The ventricle E9.5 subpopulations were named “Mt-Atp8 - CM” and “Myl3 - CM”; left ventricle E11.5 were named “Pln - CM” and “Tpm1 - CM”; right ventricle E14.5 as “Hba-a1 - CM” and “Tnnt2 - CM”; and the left atrium P0 were named “Mth6 - CM” and “Sln - CM for P0”. All of these genes used to name the cell subpopulations have been described in the literature as involved in multiple mitochondrial and cardiovascular key processes and diseases. The blue and red colors are related to down- and up-regulated transcripts, respectively, relative to one another; the numbers are related to differentially expressed transcripts in each condition (down- and up-regulated); and the numbers in parenthesis are the number of differentially expressed genes, when presenting fold-change and p-value cut-offs of 1.5 and 0.05, respectively. We observed that in all four time points, one cardiomyocyte sub-population tended to dominate with a higher number of positively expressed gene transcripts compared to another for both the coding and lncRNA transcripts, with the exception of lncRNAs P0 which had almost the same abundance of transcripts.

Next, we evaluated potential pathways and biological processes associated with the differentially expressed transcripts we identified in each cardiomyocyte subpopulation (Fig. 6 and Supplementary File 3). In the E9.5 ventricle, identified cell populations manifested associations with biological processes related mainly to RNA processing and regulation (Fig. 6A). Interestingly, “Mt-Atp8-M” cardiomyocytes manifested enrichment in transcripts related to cardiomyopathies and neurological disorders known to be associated with cardiomyopathies (Leigh disease) and heart defects (Pitt-Hopkins syndrome). On the other hand, the “Myl3-CM” cardiomyocytes are enriched in transcripts associated with muscular diseases such as spinal muscular atrophy, myelofibrosis, and anterior ischemic optic neuropathy (Fig. 6B).

**Fig. 6.**
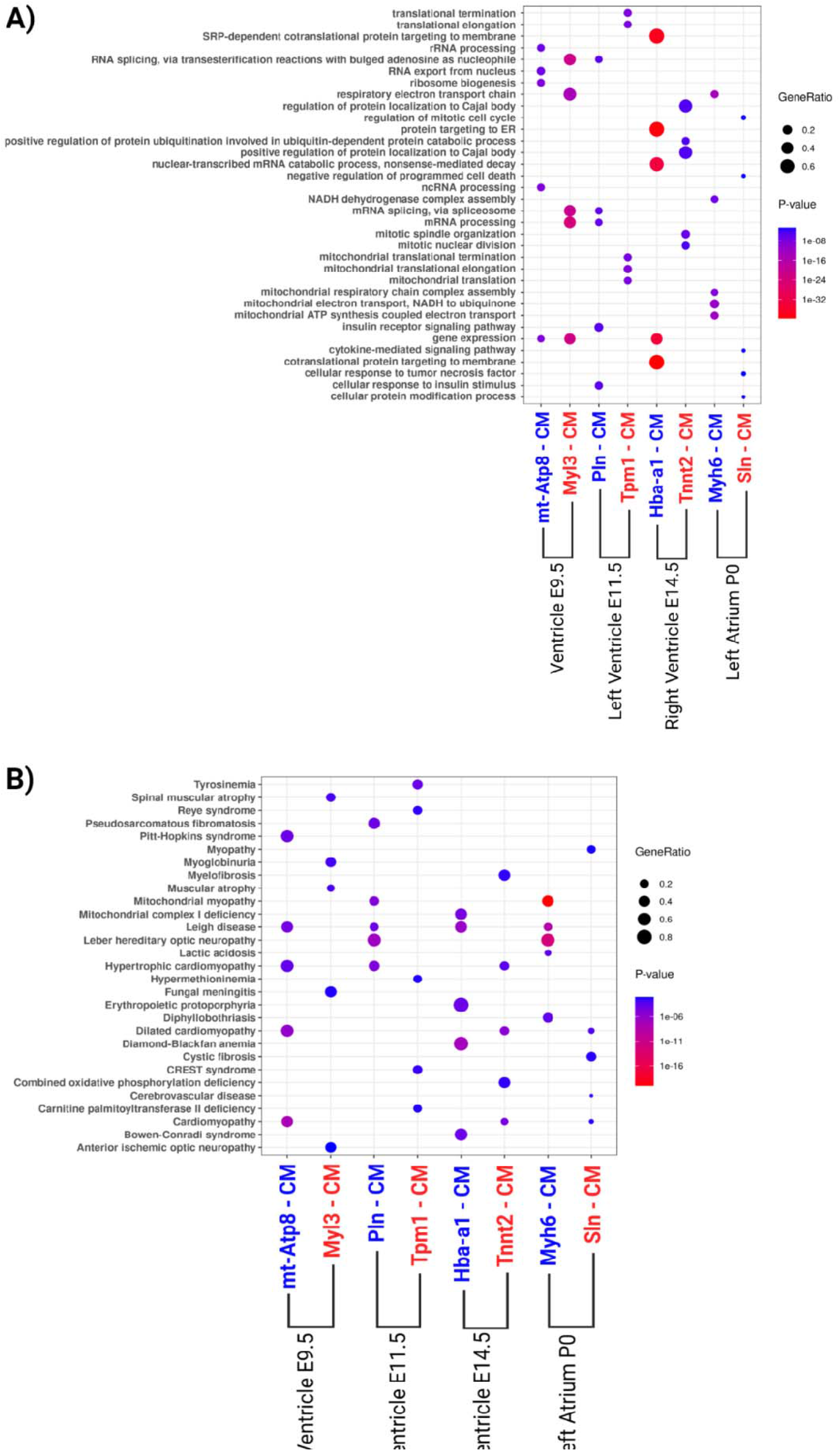
Enrichment of biological processes based on Gene Ontology (GO) pathway analysis and diseases in which the differential expressed genes are involved. The gene ratio size was computed through the number of overlap (observed/expected) genes. We named the cardiomyocyte’ populations according to the coding gene with higher expression in each chamber and time point. **A)** Biological processes from GO analysis of each cardiomyocyte sub-population. **B)** Diseases enrichment analysis of each cardiomyocyte sub-population according to Jansen Disease Database. The blue and red colors are related to down- and up-regulated transcripts, respectively.

Regarding the left ventricle E11.5, the “Pln-CM” subpopulation was similarly associated with biological processes related to RNA processing, but also with gene ontology terms linked to the insulin receptor signaling pathway (Fig. 6A). It was also enriched with genes known to be related to hypertrophic and mitochondrial myopathies, pseudosarcomatous fibromatosis, Leigh disease, and Leber hereditary optic neuropathy (Fig. 6B). On the other hand, the “Tpm1-CM” sub-population is enriched with transcripts associated with mitochondrial translational termination and elongation (Fig. 6A), and with different metabolic disorders such as tyrosinemia, Reye syndrome, hypermethioninemia, CREST syndrome, and carnitine palmitoyltransferase II deficiency (Fig. 6B).

In the right ventricle, the E14.5 “Hba-a1-CM” subpopulation was enriched with genes related to protein targeting the membrane and the endoplasmic reticulum (Fig. 6A). They were also associated with Leigh disease, as well as with blood-related diseases (erythropoietic protoporphyria and Diamond-Blackfan anemia), mitochondrial complex I deficiency, and ribosome-related disorder called Bowen-Conradi syndrome. “Tnnt2-CM” cardiomyocytes manifested enrichment in genes associated with the regulation of protein localization to the Cajal body, regulation of protein ubiquitination, mitotic spindle organization, and nuclear division. Additionally, they are associated with myelofibrosis, cardiomyopathies, and combined oxidative phosphorylation deficiency (Fig. 6B).

Finally, P0 left atrium cardiomyocytes “Myh6-CM’’ manifested enrichment in genes associated with mitochondrial function and respiratory electron transport chain (Fig. 6A), with a direct link with mitochondria-related diseases, such as mitochondrial myopathy, as well as with Leigh disease, Leber hereditary optic neuropathy, lactic acidosis, and diphyllobothriasis (Fig. 6B). On the other hand, the “Sln - CM” sub-population was enriched with genes associated with the cell cycle, protein modification process, and cytokine-mediated signaling pathway (Fig. 6A). They were also linked with cardiomyopathies, besides cystic fibrosis, and cerebrovascular disease (Fig. 6B). Fig. 6 represents the Gene Ontology (GO) Biological Processes and Jensen Diseases database enrichment analysis for each cardiomyocyte sub-populations top 5 terms. The remaining enriched gene ontology terms and categories, as well as KEGG pathways and Jensen diseases for all cardiomyocyte populations were made available in Supplementary File 3.

### Modular expression analysis reveals cell-specific functional insights for lncRNAs during myocardial development

Transcriptional modules based on co-expression patterns can shed light on novel functional connections between lncRNAs and mRNAs [25] during heart development in a single-cell perspective. In summary, identified modules can be used to redefine cell populations, reveal novel gene associations, and predict gene function by guilt-by-association. We employed CEMiTool to identify expression modules at each time point, derive gene ontology terms over-represented in each module, retrieve protein-protein and co-expression interactions within module transcripts and perform cell-specific gene set enrichment analysis (GSEA) [26]. CEMiTool was able to generate modules for 13 out of 15-time points and heart compartments, without being able to retrieve modular patterns based on guilt-by-association for atrium E9.5 and left ventricle E14.5 (Supplementary File 3).

Identified modules were differentially enriched in distinct cell types and functional processes (Fig. 7A-C). Our examination revealed modules related to cardiac tissue development (*e*.*g*. “muscle structure development”, “trabeculae formation”, “cardiac muscle cell differentiation”, “cardiac cell development”, “blood vessel morphogenesis”, “angiogenesis”, “vasculogenesis”, “vasculature development”, “ventricular cardiac muscle cell differentiation”), mitochondrial processes (*e*.*g*. “mitochondrial matrix”, “mitochondrial protein complex”, “mitochondrial translation”), cardiac functioning (*e*.*g*. “heart process”, “muscle contraction”, “cardiac conduction”, “response to cadmium ion”) and circulatory function (*e*.*g*. “platelet alpha granule”, “response to wounding”, “circulatory system process’’) (Fig. 7A). For instance, the Module 1 (M1) from the left ventricle E11.5 is enriched with coding genes known to be associated with mitochondrial processes, and the GSEA analysis shows a clear difference in the module expression between cardiomyocyte subpopulations Pln-CM and Tpm1-CM (Fig. 7B). GSEAs were undertaken separately for each time point and heart compartment, revealing cell-specific expression patterns for some of these modules (Fig. 7C). This latter analysis revealed modules harboring different expression patterns in cardiomyocyte and macrophage subpopulations (Supplementary File 4).

**Fig. 7.**
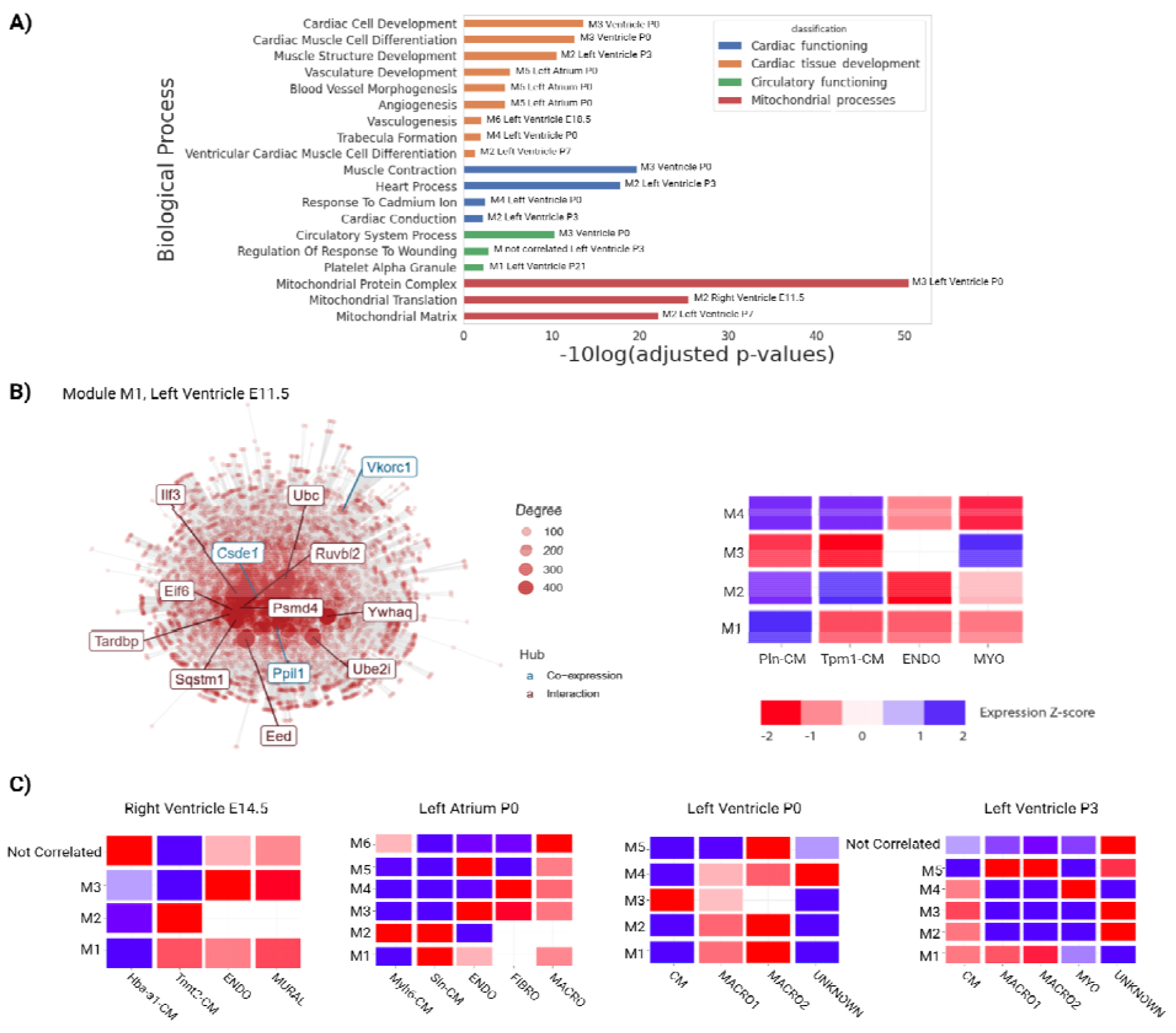
Modular expression analysis reveals cell-specific functional insights for lncRNAs along with heart development. **A)** Biological processes from GO analysis of the modules that are related to the development of cardiac tissue, mitochondrial processes, cardiac functioning, and circulatory functioning. We observe the compartment and time point to which the adjusted p-value corresponds. **B)** Module 1 (M1) from the Left ventricle E11.5 is enriched in mitochondrial processes and GSEA shows a difference between cardiomyocyte subpopulations 1 and 2. The co-expression network shows the gene’s interaction based on their expression patterns and protein-protein interaction, the gene’s names that appear are the hub genes that have more connectivity with others. **C)** GSEAs were undertaken separately for each time point and heart compartment, revealing a cell-specific expression pattern for some of these modules. The blue and red colors are related to down- and up-regulated transcripts, respectively.

In addition, selected enrichment analysis and modules were pursued which are directly related to the development of cardiac tissue (Fig. 8) as follows: in Fig. 8A we had right ventricle samples from E14.5, and we can observe mitochondrial module with 2 cardiomyocyte sub-populations, in which, these 2 had different gene set enrichment analysis (*i*.*e*. Hba-a1-CM is under-represented and Tnnt2-CM is over-represented); Fig. 8B highlights some lncRNAs which compound the module available in Fig. 8A. Some of these lncRNAs have been reported previously as directly related to heart processes (*Mhrt, Tln1, Hand2os1, Dancr, Mbnl1, Meg3, TCONS_00151380, AA465934, Snhg5*, and *1700021F05Rik*) [2,27– 35]. In addition, *Snhg5* is classified as a pcRNA. Otherwise, also exists some lncRNAs which have little expression in some module’s cell types (*AC144860*.*1, Gm15614, Gm42639, Gm42856, Gm47585, TCONS_00041212*, and *TCONS_00184519*). *Mhrt* and *Dancr* are contained in both cases: related to heart processes and have a little expression in some module’s cell types. Fig. 8C presents the module’s enrichment analysis according to Gene Ontology Biological Processes.

**Fig. 8.**
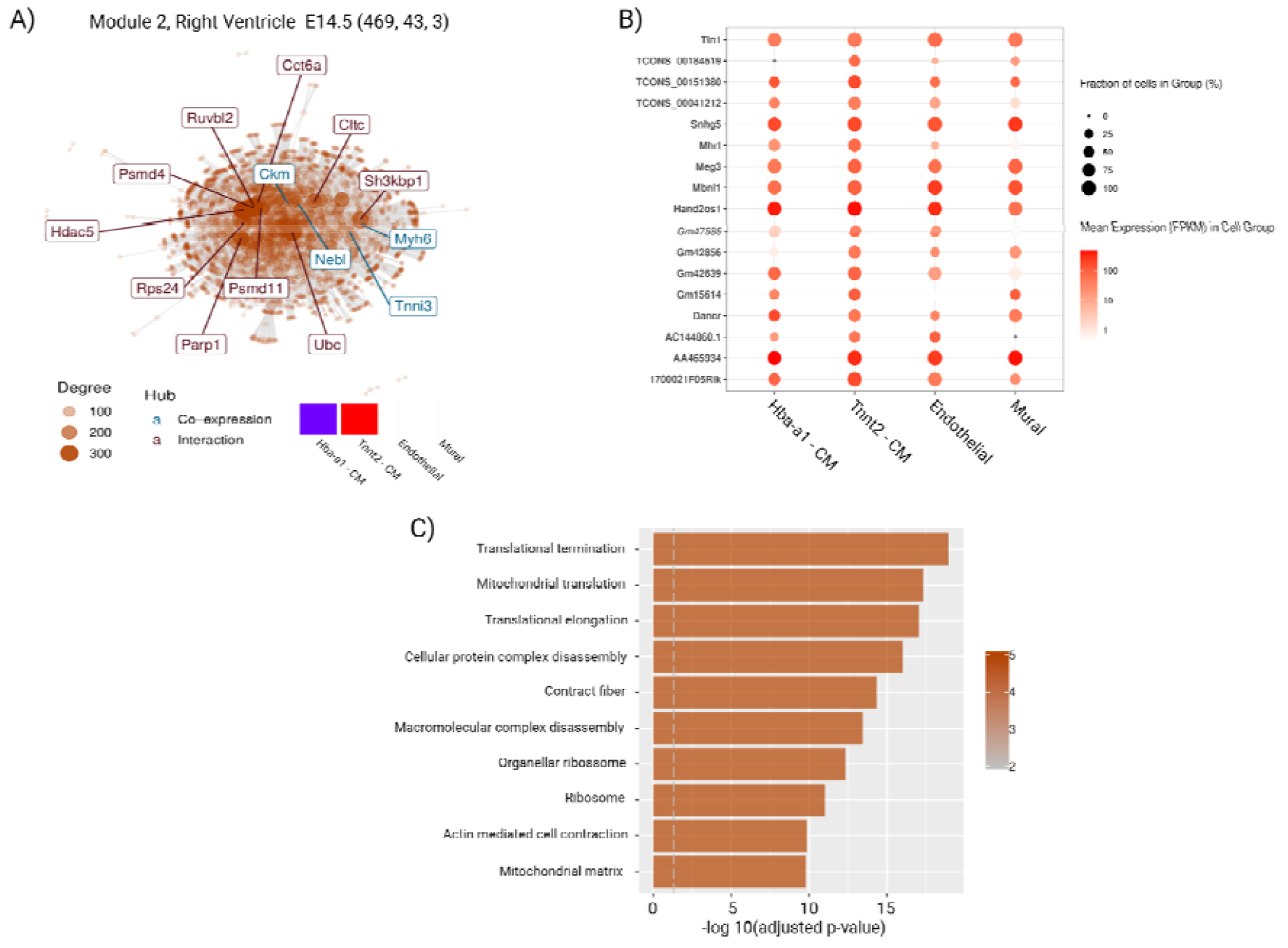
Expression of lncRNAs available in Module 2 from right ventricle E14.5. **A)** Co-expression and protein-protein interaction network from Module 2, right ventricle samples from E14.5. The small heatmap shows the GSEA analysis. The numbers 469, 43 and 3 are related to the number of coding, lncRNAs and pcRNAs that compounds in this module. **B)** The cell’s fraction and FPKM mean expression of 17 selected lncRNAs which compound the module. **C)** Represents the top 10 GO terms based on the module’s enrichment analysis.

As depicted in Fig. 8, we observed *Tln1* FPKM expression means of 42.7 and 47.55 in Hba-a1-CM and Tnnt2-CM, respectively, whereas in endothelial cells we observed expression at 70.72. *Tln1* is expressed at low levels in normal CMs compared with other cell types (*e*.*g*., in endothelial cells). In addition, we found a *Snhg5* FPKM expression mean of 265.8 in mural cells, greater than the other cell types, this pcRNA contributes to angiogenesis in acute myelogenous leukemia (Supplementary File 4).

Thus, it was noted that some stages and compartiments (ventricle E9.5, left ventricle E11.5, right ventricle E14.5 and left atrium P0) manifesting cardiomyocyte subpopulations. Regarding co-expression analysis, t-sne plot and the heatmaps, it was able to see that these subpopulations had different behaviors.Thus, it was noted that in stages of myocardial development and tissue compartments manifesting cardiomyocyte subpopulations, it was logical to separate co-expressive differences between them (ventricle E9.5, left ventricle E11.5, right ventricle E14.5 and left atrium P0), in addition to the t-SNE and heatmaps, evidencing different behaviors (*e*.*g*. Fig. 2, Supplementary Fig. 1). Likewise, clusters classified as unknown (left ventricle P0 and left ventricle P3) manifested unique co-expression behaviors compared to other types in the same condition, Thus, these cells would represent another cell type for which we do have specific markers. Of note, we cannot attribute a cell type based on these top markers defined by the M3Drop package because it identifies only 1 marker for these unknown cells (Qk - left ventricle P0; and ENSMUST00000118613.6 - left ventricle P3).

Moreover, cardiomyocyte subpopulations have been described previously in the literature [36], as well as lncRNAs associated with these subpopulations [36,37]. In fact, we can point to *Gas5* and *Sghrt* (previously annotated as *1810058i24Rik*) that are related with cardiac hypertrophy in a rat model [37], as well as regulatory lncRNAs that appear to arrest the cell cycle [36]. In addition, we also detected that these two genes are differentially expressed lncRNAs throughout heart development and cardiomyocyte sub-populations (Supplementary File 2).

## Discussion

Gene expression is a fundamental mechanism for transmitting genetic information [38]. In an effort to dissect their myriad functions, gene expression encyclopedias catalog genes that are co-expressed in different cell types, facilitating the understanding of their functions and regulatory processes [6]. There are several types of non-coding RNAs. LncRNAs comprise the most representative transcriptional units of the mammalian genome and are associated with organ development and certain diseases [9]. The study of heart development is of significance given the toll taken by cardiovascular diseases around the world.

Based on the results presented here, it was possible to identify eight different cell types, more than that described in the literature in individual cell sequencing studies [17], from different regions and stages of heart development. Interestingly, the fibroblasts and myofibroblasts transcriptional signatures were detected at E14.5, the stage at which fibroblasts from the epicardium are known to undergo EMT into the myocardium [22]. Cell populations with monocytes and macrophages profiles appeared only at late embryonic and postnatal stages, consistent with evidence in the literature of monocyte-derived macrophages increasing in the myocardium post-birth [23].

New marker transcripts distributed in coding genes and different types of lncRNAs were identified, in addition, some found specifically in certain compartments, such as *Myl3* that is known as a ventricular light chain myosin with increased expression in slow-twitch skeletal muscle fibers; and *Sln*, which is restricted to the atrial lineage during mouse heart development. Other transcripts are shared between compartments such as *Tpm1* with established roles in cardiac looping, atrial septation, and ventricular trabeculae formation.

Different cardiomyocyte cell types were identified in distinct chambers and time points, with the ventricle and left atrium populations presenting greater differences within both cardiomyocyte types. We named the cardiomyocyte cell populations according to the coding gene with higher expression in each chamber and time point. Of note, some of these genes are associated with heart diseases, *Mt-Atp8*, for example, is enriched in cardiomyocytes and is involved in many diseases in the ventricle (*e*.*g*. hypertrophic cardiomyopathy, left ventricular hypertrabeculation syndrome (LVHT), dilated cardiomyopathy), and neuropathies [39]. *Myl3* was classified as a cardiac protein that was abundantly expressed in the heart (>260 TPM) with mosaic patterns of expression in skeletal muscle [40]. In addition, *Myl3*, known as a ventricular light chain myosin with increased expression in slow-twitch skeletal muscle fibers, [40] is robustly and homogeneously present in ventricular cardiomyocytes. *Myl3* was described as differentially expressed in the ventricle and is associated with muscle atrophy and neuromuscular diseases [41]. A study in a mouse model of Huntington’s disease also reported elevated serum levels of *Myl3* concomitant with skeletal muscle atrophy [42].

*Pln*, overexpressed in the left ventricle and called cardiac phospholamban, modulates calcium re-uptake during muscle relaxation and plays an essential role in calcium homeostasis in the heart [43]. Moreover, mutations in Pln have been associated with dilated cardiomyopathy (DCM), hypertrophic cardiomyopathy (HCM) [44], and arrhythmogenic right ventricular cardiomyopathy (ARVC) [44]. Phenotypic analysis of Tpm1 in embryos revealed roles for Tpm1 in cardiac looping, atrial septation, and ventricular trabeculae formation. Moreover, Tpm1 plays multiple roles in heart development and the formation of congenital heart defects [45]. Mutations in Tpm1 have been associated with both HCM, DCM [45,46], and with peripartum cardiomyopathy (PPCM) [46].

*Hba−a1* is up-regulated in response to chronic stress along with genes associated with the vascular system. A reduced expression of *Hba−a1* is associated with a neuroprotective role [47]. *Tnnt2* is a cardiac muscle troponin T that is expressed throughout heart development and in the post-natal heart, and also in the left and right ventricle [48]. *Tnnt2* is also associated with hypertrophic cardiomyopathy, dilated cardiomyopathy, and restrictive cardiomyopathies [48]. *Myh6* is a cardiac muscle myosin that is preferentially expressed in the atrial chambers and its mutation causes atrial septal defect [49]. *Myh6* was described in differentiation in adult cardiac precursor cells (CPCs), which were characterized by up-regulation of *Myh6* and, therefore, massively differentiated into cardiomyocytes [50]. Studies have also reported additional mutations in *Myh6* linked to both hypertrophic cardiomyopathy and dilated cardiomyopathy [51].

The expression of *Sln* is restricted to the atrial lineage during mouse heart development, and this pattern is conserved in other mammals, including humans [52]. *Sln* is involved with calcium ion (Ca2+) handling that plays a crucial role in the contraction and relaxation of cardiomyocytes [52]. Moreover, *Sln* is involved with dilated cardiomyopathy [53], cardiomyopathy [53], and heart failure [52].

Transcriptional gene modules based on the co-expression patterns can shed light on novel functional connections between lncRNAs and mRNAs. Systems biology analyses identified co-expression modules linked to heart development and cardiovascular disease. Our examination revealed modules related to the development of cardiac tissue, mitochondrial processes, cardiac functioning, and circulatory functioning. Our cardiomyocyte populations were enriched in mitochondrial functions, but we also identified differences between cardiomyocyte subpopulations according to Gene Set Enrichment Analysis. In addition, we identified some lncRNAs that are already described in the literature as lncRNAs directly related to heart processes: *Mhrt* is involved with cardiac hypertrophy and heart failure [27,28], and also protects cardiomyocytes against H2O2-induced apoptosis [29]; *Tln1* is expressed at low levels in normal CMs compared with other cell types (*e*.*g*., in endothelial cells), and combined deletion of CM *Tln1* and *Tln2* destabilized the myocardium, leading to heart failure [30]. Fig. 8 demonstrates a *Tln1* FPKM expression mean of 42.7 and 47.55 in Hba-a1-CM and Tnnt2-CM, respectively, while in endothelial cells, we observed expression at 70.72, evidencing a cell-specific higher expression pattern.

*Hand2os1* orchestrates heart development; its locus dampens *Hand2* expression to restrain cardiomyocyte proliferation, thereby orchestrating balanced development of cardiac cell lineages [31]; *Dancr* is a lncRNA that has coding potential in the human heart encoding putative micropeptides and is associated with cardiovascular diseases [32]; *Mbnl1* regulates isoproterenol□induced myocardial remodeling in vitro and in vivo [54]; *Meg3* prevents cardiac fibrosis and diastolic dysfunction [2,54]; *AA465934* is associated with early diabetic cardiomyopathy [33]; *1700021f05rik*, also known as *Mtres1* is involved in characterization of the human heart mitochondrial proteome [34]. Finally, *Snhg5* upregulation induced by YY1 contributes to angiogenesis in acute myelogenous leukemia [55]. We observed a *Snhg5* FPKM expression mean of 265.8 in mural cells, greater than the other cell types (Fig. 8). Mural cells are integral components of brain blood vessels that play important roles in vascular formation, regulation of regional cerebral blood flow, regulation of vascular stability, and homeostasis [56]. *Snhg5* has also been reported to participate in the occurrence and development of glioma [35].

Our results evidence the role of particular lncRNAs in heart development, and highlights the usage of co-expression modular approaches in the cell-type functional definition. Furthermore, the knowledge generated here unveil potential processes guided by these lncRNAs during development, which can lead to the development of improved therapeutics for cardiovascular diseases targeting genes potentially involved with the re-expression of the fetal gene expression program.

## Methods

### Datasets selection and filtering criteria

For this work, we used a cardiac single-cell RNA-seq dataset from *Mus musculus* model organism provided by DeLaughter *et al*. [17]. The dataset was generated from 1,695 single-cells from 4 embryonic (E9.5, E11.5, E14.5, E18.5) and 4 post-natal (P0, P3, P7, P21) stages, according to 5 different regions of the heart (ventricle, atrium, left ventricle, right ventricle, and left atrium), captured using the Fluidigm integrated fluidic circuits (IFC). For the lncRNA reference database, we combined lncRNAs from Gencode (M20), Ensembl (GRCm38.95) and Amaral *et al*. (2018) (*mm10CombinedNCTranscripts*.*bed*) [6] which contained 18,339, 9,074 and 15,757 lncRNAs, respectively. For coding transcripts, we selected the Gencode (M20) dataset composed of 57,966 coding transcripts. Since the lncRNAs reference dataset was built from 3 different databases, preprocessing was performed to remove the redundant sequences and generate the final dataset. In this process, we eliminated the lncRNAs with more than 50% of overlap [6] from one database to another, resulting in the final dataset of 21,044 lncRNAs. To obtain expression levels of lncRNAs and coding transcripts using the heart development database from DeLaughter *et al*. [17], we performed the following pipeline: Fastqc version 0.11.9, to execute the quality control [57]; Trimmomatic version 0.39, to trim and crop the bases with low quality as well as to remove the adapters [58], using the following parameters: 2:30:10 LEADING:5 TRAILING:5 SLIDINGWINDOW:4:28 MINLEN:32; Hisat2 version 2.1.0, to mapping next-generation sequencing reads [59], using --dta-cufflinks parameter to makes hisat2 report alignment tailored specifically for Cufflinks; Samtools version 1.3.1, to convert SAM to BAM and sort the BAM’s files [60]; Cuffnorm which is a part of Cufflinks package version 2.2.1, to report expression levels normalized by FPKM [61]. To determine expressed transcripts and its filtering to avoid sparsity issues, we used a filter of an expression value (FPKM > 0.001) [6] and transcript expression percentage in cells (transcript had to be expressed in at least 5% of the cells) [62,63]. Then, we performed transcriptome saturation at a random stage of the heart development (left ventricle at E18.5) to show that lncRNAs are identified with this pipeline (Supplementary Fig. 2).

### Machine learning clustering methodology

After this, we utilized the M3Drop R package [64], which fits a Michaelis-Menten model to the pattern of dropouts in single-cell RNA Seq data. This model was used to identify significantly variable (*i*.*e*. differentially expressed) genes for use in downstream analysis, such as clustering cells. Next, we utilized the Silhouette method [65] to estimate the best clusters and performed unsupervised hierarchical clustering with Ward linkage from scikit-learn Python library to group the cells. After this, we performed from scikit-learn Python library Principal Component Analysis (PCA) to reduce the dimensionality and t-distributed Stochastic Neighbor Embedding (t-SNE), a machine learning algorithm that defines a similar probability distribution for better visualization of the clusters with high dimensionality.

### Gene markers, cell types identification, and clustering assignment

To generate the cell marker database, we utilized cell marker gene sets from DeLaughter *et al*. [17], Gladka *et al*. [18], Farbehi *et al*. [19], and Franzén *et al*. [20], then eliminated markers that were classified for more than one cell type. The number of final gene markers for each cell type was: cardiomyocytes (70); endothelial cells (100); fibroblasts and myofibroblasts (140); macrophages and monocytes (128); B-cells (76); dendritic cells (42); glial cells (2); natural killer cells (35); T-cells (59); mural cells, vascular smooth muscle cells and pericytes (42) and myoblasts (16).

Using this cell marker unified database, we checked the frequency of appearance of these marker transcripts in cardiac single cell samples from DeLaughter *et al*. [17] and we saw that there was no significant appearance of markers from glial cells and natural killer cells; therefore, we take out these 2 cell types. Our final dataset was composed of markers for 9 cell types: cardiomyocytes; endothelial; fibroblasts and myofibroblasts; macrophages and monocytes; B-cells; dendritic; T-cells; mural cells; vascular smooth muscle cells and pericytes; and myoblasts.

We used these gene markers to assign cell types to the clusters. M3Drop [64] was used to find the marker transcripts of each cluster and chi-squared and adherence tests were used to determine the cluster’s assignment signification (*i*.*e*. p-value < 0.05). As a result, we identified 8 cell types in our dataset.

### Co-expression modules analysis with CEMiTool

In order to get the knowledge of the genes that are co-expressed and to understand the biological processes linked, we used CEMiTool [26] which is a systems biology method that identifies co-expression gene modules. Each node of the co-expression modules corresponds to a gene, and a pair of nodes is connected with an edge if there is a significant co-expression relationship between them. Using CEMiTool, we also performed comprehensive modular analyses, including the following: plot the expression profile of individual genes on each module; determine the expression activity of modules in a group of samples and cell types; perform functional enrichment analysis of modules (using Supplementary Dataset 1), and integrate co-expression results with protein-protein interaction data (using Supplementary Dataset 2). In addition, we performed a gene ontology analysis for functional categories of protein-coding genes co-expressed with these lncRNAs and then, we explored the lncRNAs which were associated with the modules that were enriched in heart development processes.

### Cardiomyocyte sub-populations differential expression and functional enrichment analysis

To identify the biological variation between cardiomyocyte sub-populations, we performed differential expression analysis among sub-populations in each time point and chamber in which two types of cardiomyocytes were identified: ventricle E9.5; left ventricle E11.5; right ventricle E14.5; and left atrium P0. Differentially expressed transcripts were determined using Cuffdiff, from Cufflinks [66]. Transcripts were considered as statistically differentially expressed when presenting fold-change and p-value cut-offs of 1.5 and 0.05, respectively. Functional overrepresentation analyses of differentially expressed transcripts were performed using different databases available in the EnrichR web tool [67], using a corrected p-value cut-off of 0.05, and considering the databases Gene Ontology Biological Process [68], KEGG pathways [69] and Jensen Diseases [70].

## Data availability

The authors declare that the data supporting the findings of this study are available within the paper and its Supplementary Information files. Any remaining data that support the results of the study will be available from the corresponding author upon reasonable request. Source data are provided with this paper

## Acknowledgements

This work was supported by grants from the NIH: HL-120732 (JAH), HL-128215 (JAH), HL-126012 (JAH), HL-147933, (JAH), HL-155765 (JAH), 14SFRN20510023 (JAH), 14SFRN20670003 (JAH), Leducq grant number 11CVD04 (JAH), Cancer Prevention and Research Institute of Texas grant RP110486P3 (JAH) and by Agencia Nacional de Investigacion y Desarrollo (ANID, Chile), FONDAP 15130011 (SL and VMC), FONDECYT 1200490 (SL) and FONDECYT 1211731 (VMC). TARR received a PhD degree fellowship from Coordenação de Aperfeiçoamento de Pessoal de Nível Superior (CAPES), Brazil.

## Author contributions

TARR conceived the project, designed and performed the experiments, conducted the analyses and wrote the manuscript. SYK validated the experiments and contributed to manuscript preparation. SL and TGG contributed to the experimental design and manuscript preparation. JAH conceived the project and contributed to manuscript preparation. VMC and TGR conceived the project, designed the experiments, supervised the research and wrote the manuscript.

## Competing interests

The authors declare no competing interests.

Correspondence and requests for materials should be addressed to T.G. do R., V. M-C or S.L.

## Notes

### Competing Interest Statement

The authors have declared no competing interest.

